# Rational design of proteins that exchange on functional timescales

**DOI:** 10.1101/113845

**Authors:** James A. Davey, Adam M. Damry, Natalie K. Goto, Roberto A. Chica

**Author notes:** These authors contributed equally to this work (James A. Davey, Adam M. Damry).

## Abstract

Proteins are intrinsically dynamic molecules that can exchange between multiple conformational states, enabling them to carry out complex molecular processes with extreme precision and efficiency. Attempts to design novel proteins with tailored functions have mostly failed to yield efficiencies matching those found in nature because standard methods do not allow for the design of exchange between necessary conformational states on a functionally-relevant timescale. Here, we develop a broadly-applicable computational method to engineer protein dynamics that we term meta-multistate design. We used this methodology to design spontaneous exchange between two novel conformations introduced into the global fold of Streptococcal protein G domain β1. The designed proteins, named DANCERs, for *Dynamic And Native Conformational ExchangeRs*, are stably folded and exchange between predicted conformational states on the millisecond timescale. The successful introduction of defined dynamics on functional timescales opens the door to new applications requiring a protein to spontaneously access multiple conformational states.

## Main Text

Proteins have found widespread application in research, industry, and medicine because they can mediate complex molecular processes with extreme precision and efficiency. Even so, continued engineering of proteins with tailored functions is essential to enable novel biotechnological applications. Computational protein design (CPD) has enjoyed considerable success in creating protein sequences that stably adopt a single targeted structure (*1–5*). However, attempts to use these methods to generate proteins that can carry out specific functions have mostly failed to match the efficiencies that are found in nature (*6–9*), suggesting that fundamental aspects of protein structure that are not currently considered in design strategies must be incorporated in order to create proteins that can approach the efficacy of naturally occurring systems. One such feature is dynamics, which have been shown to be essential for many complex protein functions (*10–13*). The development of a general strategy for the rational design of protein sequences displaying predictable dynamic properties has great potential to expand the range and functionality of designed proteins, paving the way to applications that are currently inaccessible using natural proteins.

The rational design of protein dynamics requires the prediction of sequences that can adopt the necessary conformational states for exchange. The recent development of multistate design (MSD) approaches applicable to large structural ensembles (*14–16*) has provided a method for the evaluation of protein sequence energies in the context of a large number of possible conformational states. Thus, MSD can in principle be used to assess the energy landscape of a target protein and identify sequences that can exchange between distinct states. However, introduction of functionally relevant conformational exchange into a stable protein fold is a difficult design problem as it requires *a priori* knowledge of the structural features of the relevant conformational states for dynamic exchange, including the endpoint structures and intermediate states that the protein must adopt as it undergoes this conformational transition, which are often unknown. In addition, the multivariable optimization of sequences across many conformational states presents a significant computational challenge, since sequences must be designed that not only satisfy stability requirements for multiple target structures, but also yield an energy profile that would allow exchange between structures to occur on a functionally relevant timescale.

Herein, we have developed a general procedure that addresses these challenges and enables the rational design of protein dynamics, which we termed *meta-MSD* (Fig. 1). *Meta-MSD* enables the evaluation of protein energy landscapes in order to predict sequences able to spontaneously exchange between specific states. Unlike standard MSD methodologies where states are defined by the user prior to calculation (e.g., target and off-target states), meta-MSD instead assigns the identity of the states based on their structural characteristics after rotamer optimization, enabling the unbiased prediction of the preferred state for each sequence, along with an evaluation of the relative energies of every state that the sequence can stably adopt. We applied this methodology to the design of sequences that adopt the global fold of Streptococcal protein G domain β1 (Gβ1) and spontaneously exchange between two conformations that have not been previously observed for this fold. The designed dynamic Gβ1 variants, termed DANCERs, for *Dynamic And Native Conformational ExchangeRs*, were shown to be stably folded and to exchange between the predicted conformational states on the millisecond timescale.

**Figure 1.**
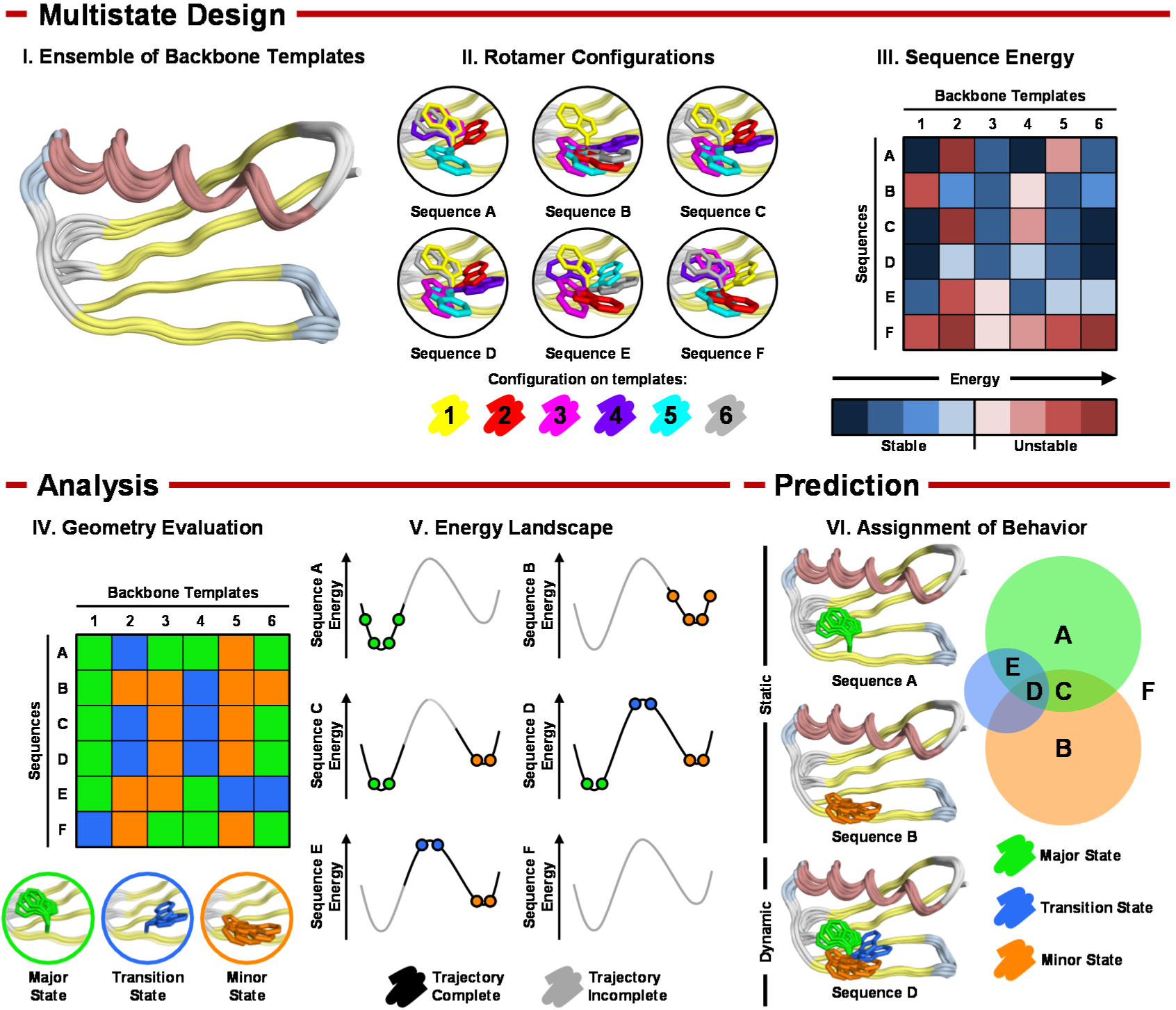
*Meta*-MSD. Multistate design (MSD) with an ensemble of backbone templates approximating the conformational landscape for dynamic exchange between targeted states (I) is used to generate microstates by solving the lowest energy rotamer configuration for each sequence on each backbone template (II). MSD also returns an energy value for each microstate that reflects its predicted stability (III). A geometry-based analysis of the rotamer-optimized microstates is performed (IV), allowing assignment of each microstate to major, minor or transition state in the energy landscape (V). Prediction of conformational dynamics is then done based on an evaluation of the relative energies of these states (VI). For a sequence to be predicted as dynamic, all three states must be stable, with an energy profile that is compatible with exchange (e.g., sequence D). Sequences A, B, and C are predicted to be static because they either stabilize a single state or cannot stabilize the transition state required for exchange. Sequence E is also predicted to be static because it stabilizes only one endpoint state. Sequence F is predicted to be unfolded because it is unstable on all states.

## Computational design of a protein energy landscape

A dynamic protein that spontaneously interconverts between two distinct conformational states adopts a continuum of unique configurations during exchange. However, the energy landscape is complex and the range of configurations that are sampled over the course of exchange cannot be completely defined. Nevertheless, it should be possible to engineer a user-defined exchange trajectory by identifying sequences that stabilize configurations having structural characteristics postulated to facilitate this exchange. To simplify the exchange reaction coordinate, the conformational landscape can be conceptually divided into three states: a major, a minor, and a transition state (Fig. S1). In the context of this work, we treat each of these states as a collection of unique configurations that we will refer to as microstates. Microstates are generated by optimizing rotamers for predefined sequences on an ensemble of backbone templates using MSD, which also returns an energy value for each microstate that reflects its predicted stability (Fig. 1, panels I-III). Following MSD, microstates are partitioned into their corresponding states according to their structural features (Fig. 1, panel IV), and the energy of each state is calculated from the energy of its constituent microstates. Evaluation of relative energies between each state then allows prediction of the exchange profile for each sequence, allowing identification of sequences that would give rise to static or dynamic Gβ1 folds (Fig. 1, panels V–VI). We call this framework *meta-* MSD because both state and dynamic behavior are assigned after rotamer optimization by MSD. *Meta-MSD* can be used to identify sequences that can stably populate the two target states, with a transition state barrier that is small enough to allow interconversion between these two states, enabling the rational design of dynamics.

To validate our meta-MSD framework, we targeted the introduction of millisecond timescale exchange into the Gβ1 structure. Native Gβ1 is rigid on this timescale (*17*), with a small size (56 amino acids) that facilitates characterization of its dynamic properties at atomic resolution. Additionally, Gβ1 possesses a single tryptophan residue (Trp43) that in high-resolution structures of Gβ1 and its natively folded variants (*18–28*) exclusively occupies a single side-chain conformation with χ1 and χ2 dihedrals of – 74 ± 9° and +75 ± 11°, respectively. We name this conformation +*g*(–) due to its positive χ2 dihedral angle and its *gauche*(*–*)χ1 dihedral (Fig. S2). In Gβ1, the Trp43 side chain is mostly solvent inaccessible, making intimate contacts with several residues that comprise the hydrophobic core. This makes it an attractive target for the design of conformational exchange, with one state being buried, and the other being excluded from the hydrophobic core in a solvent-exposed conformation that should be straightforward to distinguish spectroscopically. In addition, exchange between a core-buried and solvent-exposed state is expected to involve the disruption of side-chain interactions that should increase the kinetic barrier separating states, while not requiring large-scale changes in backbone structure that could prove kinetically inaccessible (*29*). Moreover, with exchange of the tryptophan side chain being set as our target for the design of dynamics, tryptophan side-chain dihedral angles provide a convenient metric for the assignment of microstates to one of the target states defined in our *meta-MSD* approach.

Using meta-MSD, we designed Gβ1 sequences that could adopt the native fold and also undergo conformational exchange between a state where the Trp43 indole is solvent-exposed [–*g*(+)] and a state where the indole is sequestered from the solvent in the hydrophobic core [–*g*(–)] (Fig. S3). Notably, we avoided selection of the native Trp43 conformation [+g(–)] for the core-buried state, since CPD has a tendency to overemphasize the stability of the native rotamer relative to non-native configurations (*30*). A final and particularly critical aspect of our conformational exchange design was the definition of an intermediate state with the Trp43 side chain in the −*t* conformation, since this state is necessary to provide a model of transiently populated microstates that are sampled along the reaction coordinate. Use of the –*t* conformation as a proxy of the transition state thus allowed estimation of kinetic barriers between states, enabling the elimination of sequences predicted to stably adopt two end-states separated by large kinetic barriers that would not exchange on functionally relevant timescales.

To ensure adequate sampling of the range of structures that may be required to accommodate the designed conformational exchange, an ensemble of 12,648 templates was prepared using a combination of several template generation procedures (Fig. S4, Table S1, and SI Text). Using this ensemble, MSD was performed to optimize rotamers for a library of 1,296 Gβ1 sequences comprising combinations of core-residue mutations (Fig. S5) that were previously reported to result in folded Gβ1 variants (*14*). MSD thus yielded >16 million microstates and corresponding energies, allowing for approximation of the accessible conformational landscape of Trp43 in the native Gβi fold.

Sequences having a Boltzmann-weighted average of MSD energies greater than that of the wild-type sequence are less likely to adopt a stable Gβ1 fold (*15*) and were therefore eliminated from the *meta-MSD* analysis. For the remaining 195 sequences, each microstate was classified as being in a core-buried [–*g*(–)], solvent-exposed [–*g*(+)], or intermediate [–*t*] state based on the χ_1_ and χ_2_ dihedrals of the Trp43 side chain. The energy of each of these states was determined for every sequence by taking the energy of the most stable microstate assigned to each state. State energies were used to construct an energy profile for each sequence (Fig. 1, panel V), enabling us to identify 35 sequences predicted to allow conformational exchange between the target core-buried and solvent-exposed conformations (SI Text), of which four were selected for experimental characterization (Table 1, DANCER proteins).

**Table 1.** Predicted and Experimental Properties of Gβ1 variants

| Protein | Mutations | Meta-MSD Predictions |  |  | Stability | Exchange <sup>c</sup> |  |  |  |
| --- | --- | --- | --- | --- | --- | --- | --- | --- | --- |
| | | Behavior <sup>a</sup> | $\Delta E_{eq}$ <sup>b</sup><br>(kcal/mol) | $\Delta E^{\ddagger}$ <sup>c</sup><br>(kcal/mol) | $\Delta G_U$ <sup>d</sup><br>(kcal/mol) | $k_{-1}$ <sup>f</sup><br>(s<sup>-1</sup>) | $k_1$ <sup>g</sup><br>(s<sup>-1</sup>) | $\Delta G^{\ddagger}$ <sup>h</sup><br>(kcal/mol) | $\Delta G_{eq}$ <sup>i</sup><br>(kcal/mol) |
| Wild type | | +g(-) | 8.6 | 15.2 | 4.1 ± 0.2 | | | | |
| DANCER-0 | Y3F/L5A/L7I/A34F/V39I | -g(+) ↔ -g(-) | 0.9 | 7.8 | 1.5 ± 0.2 | | | | |
| DANCER-1 | Y3F/L5A/L7I/A34F/V39L/V54I | -g(-) ↔ -g(+) | 3.7 | 8.4 | 2.2 ± 0.1 | 30 ± 10 | 110 ± 50 | 18.9 ± 0.3 | 0.3 ± 0.1 |
| DANCER-2 | Y3F/L5A/L7I/A34F/V39L | -g(+) ↔ -g(-) | 1.3 | 9.4 | 1.7 ± 0.1 | j | j | j | 1.4 ± 0.7 |
| DANCER-3 | Y3F/L7I/A34F/V39L/V54I | -g(-) ↔ -g(+) | 2.9 | 13.7 | 2.0 ± 0.3 | 3.9 ± 0.2 | 23 ± 5 | 20.65 ± 0.08 | 1.3 ± 0.3 |
| NERD-S | Y3F/L7I/A34F/V39I/V54I | -g(+) | 4.3 | 14.7 | 2.7 ± 0.1 | | | | |
| NERD-C | Y3F/L7I/F30L/V39I | +g(-) | 12.2 | 15.3 | 4.0 ± 0.3 | | | | |

<sup>a</sup> Static variants (NERD-S and NERD-C) are predicted to occupy a single state while DANCER proteins are predicted to exchange between major and minor states (major state ↔
minor state)
<sup>b</sup> Energy difference between the two lowest energy states
<sup>c</sup> Energy barrier to conformational exchange (see SI text for more detail)
<sup>d</sup> Free energy of unfolding determined by chemical denaturation with guanidium chloride at 25 °C
<sup>e</sup> Kinetic parameters ( $k_1$ , $k_{-1}$ , $\Delta G^{\ddagger}$ ) reported at 15 °C, $\Delta G_{eq}$ at 25 °C.
<sup>f</sup> Rate constant for exchange from major to minor state
<sup>g</sup> Rate constant for exchange from minor to major state
<sup>h</sup> Energy barrier for exchange from major to minor state
<sup>i</sup> Free energy difference between major and minor states

## Experimental characterization

Although the four DANCER proteins each contained between five and six mutations, representing approximately 10% of the Gβ1 total sequence length, they expressed as soluble monomers (Fig. S6), adopted the native Gβ1 fold (Fig. S7), and were folded at room temperature (Fig. S8, Table S2). Chemical denaturation experiments (Fig. S9) could be fit to a two-state model with *m*-values similar to that of the wild type (Table S2), indicating a similar level of protein surface exposed to solvent upon unfolding (*31*). In addition, all DANCER variants have unfolding free energies that are 1.5 kcal/mol and higher (Table 1), confirming that they are stably folded at room temperature. Solution NMR was used to assess the dynamic properties of DANCER proteins, with ^1^H-^15^N heteronuclear single quantum coherence (HSQC) spectra showing immediate evidence that DANCER proteins exist in two distinct conformational states (Fig. S10). Specifically, spectra for DANCER-1, DANCER-2, and DANCER-3 all showed the presence of a minor species not seen in spectra of wild-type Gβ1 (Fig. S11). The only exception was DANCER-0, which instead showed significant peak broadening, suggesting that it is dynamic on a faster timescale (*32*).

Using ^1^H-^15^N HSQC ZZ-exchange experiments (Fig. 2 and S12), we confirmed that the minor species in DANCER-1, DANCER-2, and DANCER-3 is an alternate state of Gβ1 undergoing exchange with the major species. Mixing-time dependent changes in peak intensities acquired over a range of temperatures could be fit to kinetic and thermodynamic parameters of exchange for DANCER-1 and DANCER-3 (Table 1, Fig. S13), confirming that conformational exchange is occurring on the millisecond timescale. DANCER-1 exhibits approximately 10-fold faster exchange than DANCER-3, with an activation barrier that is 1.75 kcal/mol smaller in magnitude. Conformational exchange was also observed for DANCER-2, although the small population of the minor state (< 10%) prevented quantitative measurement of kinetic parameters for this mutant.

**Figure 2.**
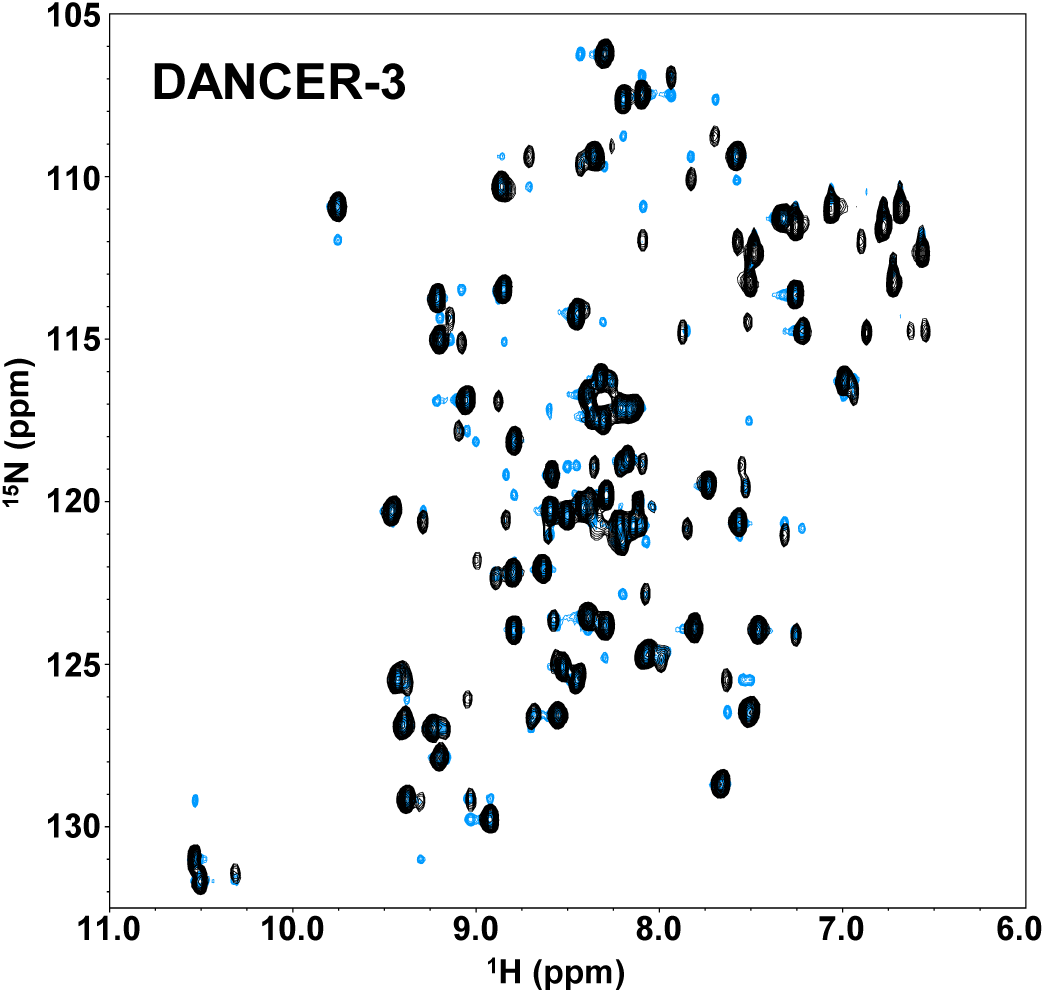
^J^H-^15^N ZZ-Exchange spectrum for DANCER-3. ZZ-Exchange spectrum (blue) is shown overlaid with ^1^H-^15^N HSQC spectrum (black) to highlight the presence of exchange peaks.

To obtain structural evidence that the two conformations sampled by our dynamic Gβ1 variants matched structural states predicted by *meta*-MSD, solution NMR was used to solve the structure of the major state of DANCER-2 (Fig. 3A, Table S3). As predicted, this structure shows a native Gβ1 fold with χ_1_ and χ_2_ dihedrals for Trp43 that correspond to the solvent-exposed –*g*(+) conformation (Table 2). However, there was also a secondary network of low intensity NOEs involving the Trp43 side chain that were not compatible with this structure, but could be used to determine a structural model for the alternate, minor state (SI Text). According to this model (Fig. 3B, Table S3), the configuration of Trp43 in the minor state is in the core-buried −g(−) state (Table 2), as predicted by *meta*-MSD. Taken together, these data demonstrate that we have successfully designed a sequence that adopts the Gβ1 fold while undergoing conformational exchange on a millisecond timescale between two conformational states that have not previously been observed, but were the targets of our design protocol.

**Figure 3.**
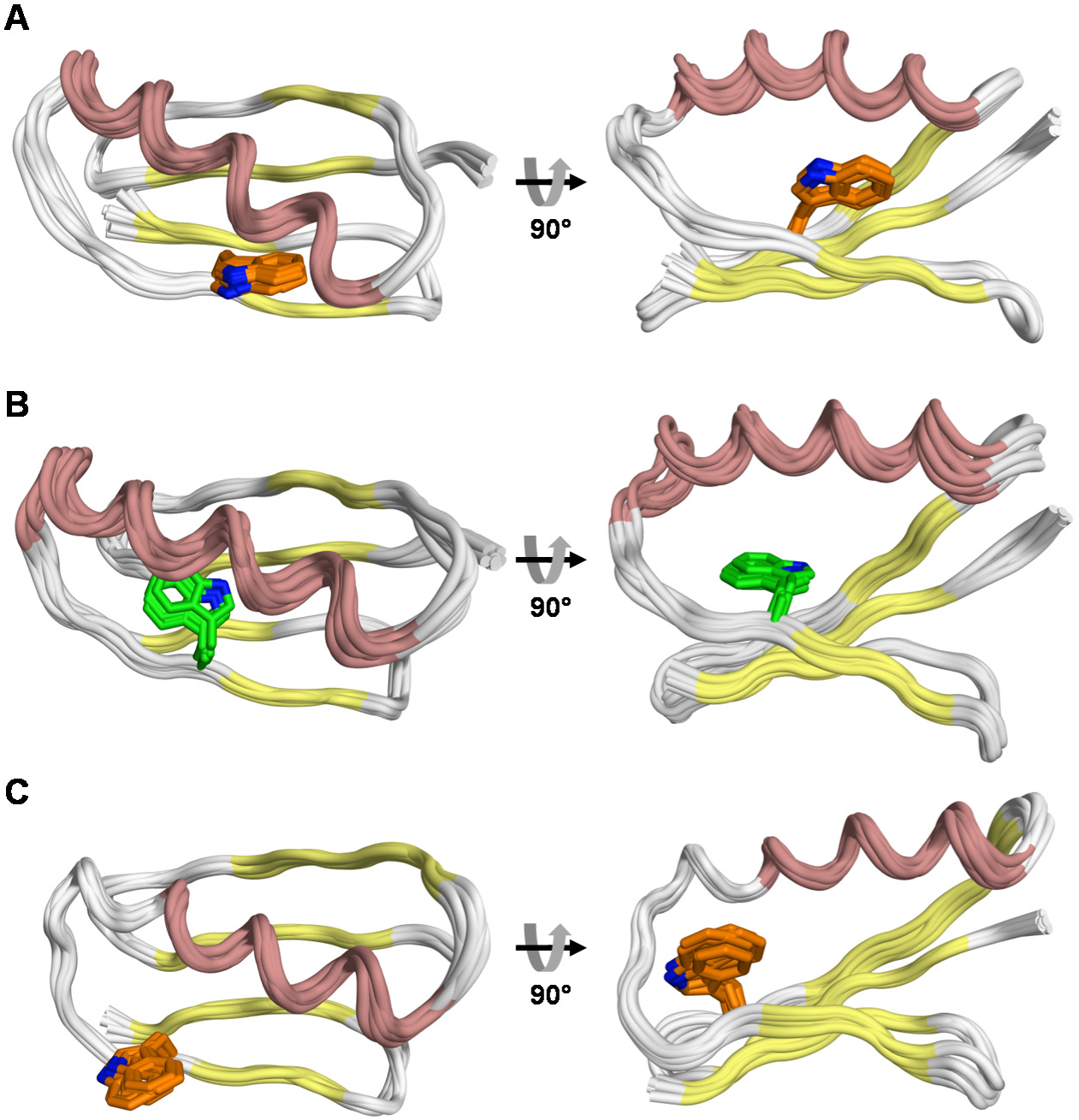
Figure 3. Solution structures of Gβ1 variants. NMR ensembles for (A) DANCER-2 major species, (B) DANCER-2 minor species, and (C) NERD-S. The minor species of DANCER-2 is a model generated using NOESY data that excluded a small subset of peaks from the automatic NOE assignment process that could be unambiguously assigned to the major species (SI Text). The Trp43 side chain is shown as sticks.

**Table 2.** Comparison of predicted and experimental structures

| Protein | TM-score to 1PGA <sup>a</sup> | Predicted<br>Trp43 Conformation | Experimental $\chi_1$<br>(°) | Experimental $\chi_2$<br>(°) |
| --- | --- | --- | --- | --- |
| **DANCER-2** |  |  |  |  |
| Major species | 0.67 | -g(+) | +75 ± 2 | -74 ± 1 |
| Minor Species | 0.66 | -g(-) | -95 ± 1 | -110 ± 2 |
| **Static Gβ1 variants** |  |  |  |  |
| NERD-S | 0.66 | -g(+) | +54 ± 4 | -89 ± 2 |
| NERD-C | 0.85 | +g(-) | -84 ± 4 | +80 ± 4 |

<sup>a</sup> TM-score has a value between 0 and 1, where 1 indicates a perfect match between two structures. Two proteins with a TM-score greater than 0.5 are considered to adopt the same fold (35, 36).

To illustrate the reliability of our meta-MSD predictions, we also characterized the structure and dynamics of DANCER-1 and DANCER-3. While the exchange parameters for these mutants made it impractical to attempt structure determination, ^1^H-^15^N HSQC spectra of the major species showed similarities with those of other structurally characterized variants, suggesting a high degree of structural similarity with these states. Specifically, the DANCER-1 spectrum shows only small chemical shift differences from that of DANCER-2 (Fig. 4A), suggesting that the major species of DANCER-1 also contains Trp43 in the solvent-exposed −g(+) state. Likewise, the ^1^H-^15^N HSQC spectrum for DANCER-3 was highly similar to that of a variant that we determined to thermodynamically and kinetically favor the −*g*(+) state as predicted by *meta*-MSD (Fig. 4B), called NERD-S, for *Non-Exchanging Rigid Design* with a *Solvent-exposed* Trp43 side chain (SI Text, Fig. 3C, Tables 1, 2, and S3). Therefore in all three of the mutants predicted by meta-MSD to be dynamic for which structural information could be obtained, the major conformation was the Gβ1 structure with Trp43 being in the solvent-exposed –*g*(+) state.

**Figure 4.**
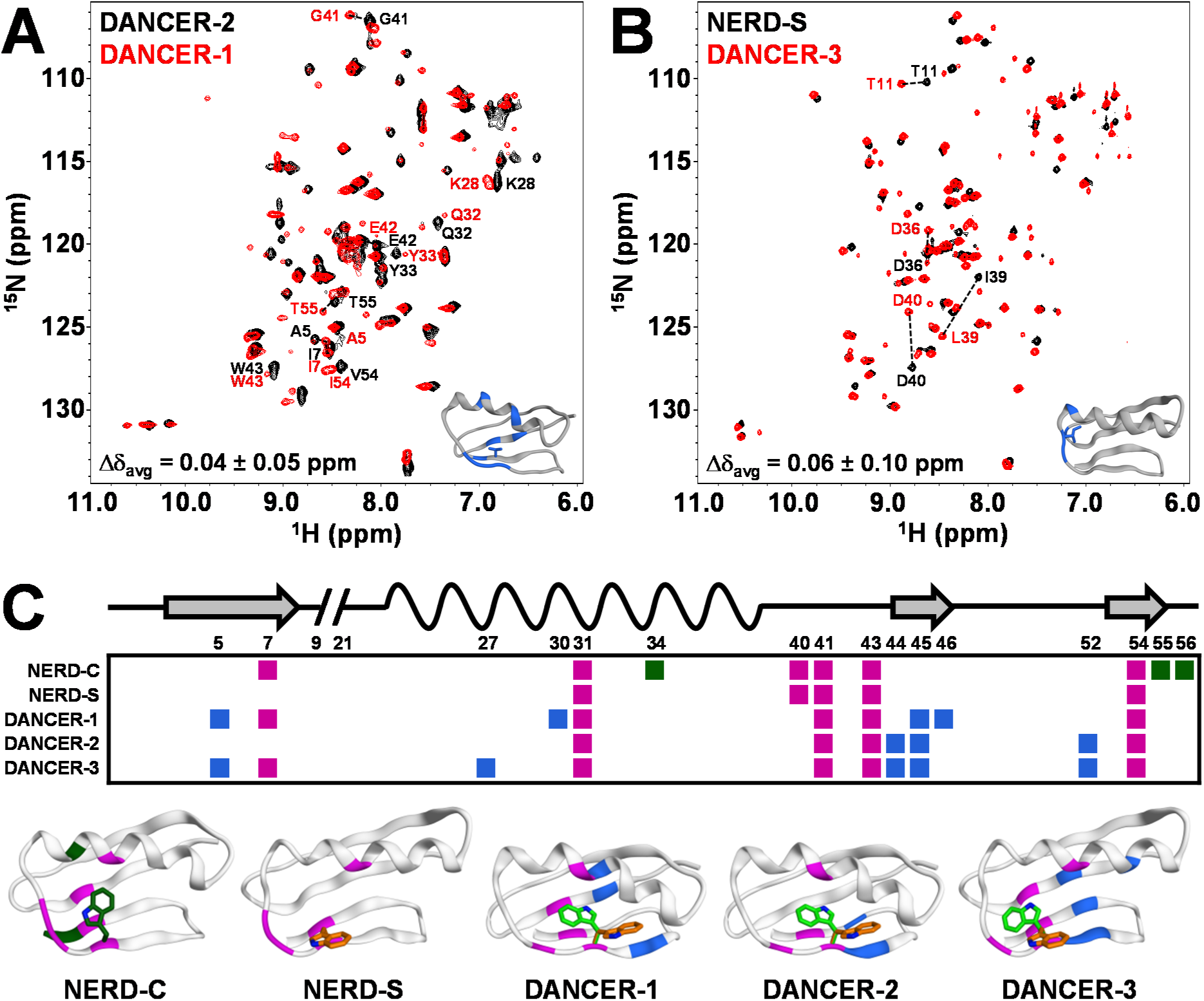
Structural analysis of DANCER-1 and DANCER-3. (A) Superimposed ^1^H-^15^N-HSQC spectra of DANCER-2 and DANCER-1 reveal high structural similarity between major states. Residues showing significant average amide shift differences (Δ δ > (Δ δ_avg_ + 1σ) are labeled and highlighted in blue on the inset DANCER-2 structure. These residues are all proximal to the single amino acid that differs between the two DANCER proteins (shown as sticks). (B) ^1^H-^15^N-HSQC spectra demonstrating that the major state of DANCER-3 has the same structure as NERD-S. (C) Summary of NOE correlations involving the Trp43 indole N-H shown on a position map (secondary structure elements on top) and on each structure. Correlations are colored green, blue, or magenta, if they are observed in static, dynamic, or both variants respectively. Trp43 side-chain conformation(s) consistent with observed NOEs are shown for each structure. Included in this analysis is the solution NMR structure of NERD-C (*Non-Exchanging Rigid Design* with a *Coreburied* Trp43 conformation, SI Text), which adopts the native +g(–) configuration (Fig. S15, Table 2). NERD-C shows several unique indole N-H NOE correlations that are not observed in any of the DANCER variants, confirming that this state is not sampled by the DANCER proteins.

Insight into the minor states being sampled in the conformational exchange exhibited by DANCER-1 and DANCER-3 was provided by ^1^H-^15^N-NOE correlations involving the Trp43 indole NH proton. DANCER-1 and DANCER-3 spectra show NOE correlations (Fig. S14) to similar regions of the protein as was observed in DANCER-2 (Fig. 4C), consistent with exchange between core-buried –*g*(−) and solvent-exposed −*g*(+) states. Furthermore, comparison of NOEs involving the Trp43 indole NH proton confirmed that these correlations do not correspond to the core-buried state found in the wild-type structure [+*g*(−)] (SI text, Fig. S15). Taken together, our NMR results confirm that the Trp43 residues of DANCER-1 and DANCER-3 exchange between the solvent-exposed −g(+) and core-buried *−g*(−) conformations that were the targets of our design, and also suggest that exchange is achieved via a coordinated change in side-chain configurations for a triad of aromatic residues (Phe34, Trp43, Phe45) in a process we have termed an aromatic relay (SI text, Fig. S16).

## Discussion

The *meta*-MSD framework described here enabled the rational design of Gβ1 variants that spontaneously exchange between two predefined states on the millisecond timescale without the need for an external stimulus to induce exchange. To our knowledge, this work represents the first successful application of CPD to engineer a specific mode of conformational exchange into a stable protein fold. Although a previous CPD-based design generated a protein capable of reversible exchange between coiled-coil trimer and zinc-finger folds (*8*), this relied on the presence of a metal that was critical for the formation of the zinc finger structure. In that case, it was possible to design exchange by simultaneously minimizing the sum of the sequence energies across both folds. In contrast, to design conformational exchange between two states in the absence of a ligand or other external stimulus, we found that it was essential to explicitly consider both the relative energies between the two target end-states (ΔEeq) and the barrier to conformational exchange (ΔE^‡^). Without estimation of both of these energy differences, it would not have been possible to distinguish between dynamic (DANCER) and static (NERD) sequences (e.g., both (ΔE_eq_ and (ΔE^‡^ values for DANCERs were lower than for NERDs and wild-type Gβ1, Table 1).

Another key advantage arising from our utilization of a *meta*-analysis-based design strategy is that it enabled the use of a significantly larger structural ensemble than has previously been utilized in MSD approaches. This was of critical importance, since we found that the full complement of seed structure and ensemble generation strategies used in our framework was required to approximate the energy landscape of the designed exchange trajectory with enough accuracy to predict DANCER variants (SI Text, Table S4). In addition, the large ensemble size made it possible to design exchange in the absence of specific structures corresponding to each end-state, in contrast with the metal-triggered conformational exchange that was previously designed using available crystal structures as templates for the two end-states (*8*).

Importantly, our results show that the introduction of dynamics on the millisecond timescale cannot be achieved via a single mutation and that instead dynamics is conferred through subtle interactions across a network of residues. For example, the A34F mutation, which was previously shown to induce dimerization of Gβ1 without altering the Trp43 conformation (*27, 33*), is common to all DANCER proteins and an integral component of the aromatic relay that underlies exchange (Fig. S16). However, this mutation alone is not sufficient to introduce dynamics into the Gβ1 fold, since the variant NERD-S also possesses this mutation but does not undergo exchange on the millisecond timescale (Table 1). Introduction of the conservative and isosteric I39L mutation into the NERD-S sequence appears to be sufficient to introduce the targeted conformational exchange, giving rise to the dynamic variant DANCER-3. These results highlight the challenges of attempting to infer dynamics from simple sequence characteristics, and demonstrate the power of *meta-MSD* to design conformational exchange into proteins even without prior knowledge of the mechanism of exchange.

## Conclusion

The *meta*-MSD framework presented here is in principle applicable to the design of specific conformational exchange into any globular protein. In the future, *meta*-MSD could also be used to design proteins with functions that rely on the ability to spontaneously access more than one conformational state (*e.g.* open and closed states of an enzyme to facilitate substrate binding and catalysis, respectively). Alternatively, *meta*-MSD could be used to enrich functionally relevant but low occupancy states from an ensemble of dynamic configurations to improve function *(34).* Moreover, while we have demonstrated the introduction of dynamics into a rigid protein, dampening of dynamics should in principle also be possible, as demonstrated by our design of NERD-C and NERD-S. This potential for *meta*-MSD to be used for the rigidification of highly dynamic regions in proteins without adversely affecting the overall structure, in effect imitating conformational selection *in silico*, opens the door to the design of proteins with a wider range of functions than previously possible.

## Data Availability

Structure coordinates have been deposited in the Protein Data Bank with accession codes 5UB0 (NERD-C), 5UBS (NERD-S), 5UCE (major state of DANCER-2), and 5UCF (minor state of DANCER-2). NMR data has been deposited in the Biological Magnetic Resonance Data Bank with accession codes 30220 (NERD-C), 30221 (NERD-S), 30222 (DANCER-2), 27030 (DANCER-0), 27031 (DANCER-1), and 27032 (DANCER-3).

## Supplementary Materials

Supplementary Text

Figs. S1 to S16

Tables S1 to S4

## Acknowledgments

R.A.C. acknowledges grants from the Natural Sciences and Engineering Research Council of Canada (NSERC), the Ontario Research Fund, and the Canada Foundation for Innovation. N.K.G. acknowledges a grant from NSERC. J.A.D. is the recipient of an Ontario Graduate Scholarship and A.M.D. is the recipient of a NSERC postgraduate scholarship. We acknowledge Dr. Glenn Facey, Dr. Yves Aubin, and Simon Sauvé for assistance with NMR experiments, as well as Dr. Yun Mou for helpful discussions.

## Author Contributions

J.A.D. and A.M.D. performed the experiments and analyzed data. N.K.G. and A.M.D. designed NMR experiments and analyzed data. J.A.D. and R.A.C. designed computational experiments. All authors wrote the manuscript.

## References

1. B. I. Dahiyat, S. L. Mayo, De novo protein design: fully automated sequence selection. Science 278, 82–87 (1997).

2. B. Kuhlman, G. Dantas, G. C. Ireton, G. Varani, B. L. Stoddard, D. Baker, Design of a novel globular protein fold with atomic-level accuracy. Science 302, 1364–1368 (2003).

3. N. Koga, R. Tatsumi-Koga, G. Liu, R. Xiao, T. B. Acton, G. T. Montelione, D. Baker, Principles for designing ideal protein structures. Nature 491, 222–227 (2012).

4. E. Marcos, B. Basanta, T. M. Chidyausiku, Y. Tang, G. Oberdorfer, G. Liu, G. V. Swapna, R. Guan, D. A. Silva, J. Dou, J. H. Pereira, R. Xiao, B. Sankaran, P. H. Zwart, G. T. Montelione, D. Baker, Principles for designing proteins with cavities formed by curved beta sheets. Science 355, 201–206 (2017).

5. S. M. Malakauskas, S. L. Mayo, Design, structure and stability of a hyperthermophilic protein variant. Nat Struct Biol 5, 470–475 (1998).

6. L. Jiang, E. A. Althoff, F. R. Clemente, L. Doyle, D. Röthlisberger, A. Zanghellini, J. L. Gallaher, J. L. Betker, F. Tanaka, C. F. Barbas 3rd,, D. Hilvert, K. N. Houk, B. L. Stoddard, D. Baker, De novo computational design of retro-aldol enzymes. Science 319, 1387–1391 (2008).

7. S. Bjelic, L. G. Nivón, N. Çelebi-Ölçüm, G. Kiss, C. F. Rosewall, H. M. Lovick, E. L. Ingalls, J. L. Gallaher, J. Seetharaman, S. Lew, G. T. Montelione, J. F. Hunt, F. E. Michael, K. N. Houk, D. Baker, Computational design of enone-binding proteins with catalytic activity for the Morita-Baylis-Hillman reaction. ACS Chem Biol 8, 749–757 (2013).

8. X. I. Ambroggio, B. Kuhlman, Computational design of a single amino acid sequence that can switch between two distinct protein folds. J Am Chem Soc 128, 1154–1161 (2006).

9. H. K. Privett, G. Kiss, T. M. Lee, R. Blomberg, R. A. Chica, L. M. Thomas, D. Hilvert, K. N. Houk, S. L. Mayo, Iterative approach to computational enzyme design. Proc Natl Acad Sci USA 109, 3790–3795 (2012).

10. G. Bhabha, J. Lee, D. C. Ekiert, J. Gam, I. A. Wilson, H. J. Dyson, S. J. Benkovic, P. E. Wright, A dynamic knockout reveals that conformational fluctuations influence the chemical step of enzyme catalysis. Science 332, 234–238 (2011).

11. J. S. Fraser, M. W. Clarkson, S. C. Degnan, R. Erion, D. Kern, T. Alber, Hidden alternative structures of proline isomerase essential for catalysis. Nature 462, 669–673 (2009).

12. S. J. Kerns, R. V. Agafonov, Y. J. Cho, F. Pontiggia, R. Otten, D. V. Pachov, S. Kutter, L. A. Phung, P. N. Murphy, V. Thai, T. Alber, M. F. Hagan, D. Kern, The energy landscape of adenylate kinase during catalysis. Nat Struct Mol Biol 22, 124–131 (2015).

13. S. R. Tzeng, C. G. Kalodimos, Dynamic activation of an allosteric regulatory protein. Nature 462, 368–372 (2009).

14. B. D. Allen, A. Nisthal, S. L. Mayo, Experimental library screening demonstrates the successful application of computational protein design to large structural ensembles. Proc Natl Acad Sci U S A 107, 19838–19843 (2010).

15. J. A. Davey, R. A. Chica, Improving the accuracy of protein stability predictions with multistate design using a variety of backbone ensembles. Proteins 82, 771–784 (2014).

16. J. A. Davey, A. M. Damry, C. K. Euler, N. K. Goto, R. A. Chica, Prediction of Stable Globular Proteins Using Negative Design with Non-native Backbone Ensembles. Structure 23, 2011–2021 (2015).

17. K. A. Crowhurst, S. L. Mayo, NMR-detected conformational exchange observed in a computationally designed variant of protein Gbeta1. Protein Eng Des Sel 21, 577–587 (2008).

18. S. Butterworth, V. Lamzin, D. Wigley, J. Derrick, K. Wilson, Anisotropic refinement of a protein G domain at 1.1 ångstrom resolution. The Protein Databank in Europe (accessed 13.11.2012). Available online at: http://www.ebi.ac.uk/pdbe, (1998).

19. J. P. Derrick, D. B. Wigley, The third IgG-binding domain from streptococcal protein G. An analysis by X-ray crystallography of the structure alone and in a complex with Fab. J Mol Biol 243, 906–918 (1994).

20. T. Gallagher, P. Alexander, P. Bryan, G. L. Gilliland, Two crystal structures of the B1 immunoglobulin-binding domain of streptococcal protein G and comparison with NMR. Biochemistry 33, 4721–4729 (1994).

21. A. M. Gronenborn, D. R. Filpula, N. Z. Essig, A. Achari, M. Whitlow, P. T. Wingfield, G. M. Clore, A novel, highly stable fold of the immunoglobulin binding domain of streptococcal protein G. Science 253, 657–661 (1991).

22. B. J. Wylie, L. J. Sperling, A. J. Nieuwkoop, W. T. Franks, E. Oldfield, C. M. Rienstra, Ultrahigh resolution protein structures using NMR chemical shift tensors. Proc Natl Acad Sci USA 108, 16974–16979 (2011).

23. J. H. Tomlinson, V. L. Green, P. J. Baker, M. P. Williamson, Structural origins of pH-dependent chemical shifts in the B1 domain of protein G. Proteins 78, 3000–3016 (2010).

24. D. J. Wilton, R. B. Tunnicliffe, Y. O. Kamatari, K. Akasaka, M. P. Williamson, Pressure-induced changes in the solution structure of the GB1 domain of protein G. Proteins 71, 1432–1440 (2008).

25. P. Strop, A. M. Marinescu, S. L. Mayo, Structure of a protein G helix variant suggests the importance of helix propensity and helix dipole interactions in protein design. Protein Sci 9, 1391–1394 (2000).

26. T. Saio, K. Ogura, M. Yokochi, Y. Kobashigawa, F. Inagaki, Two-point anchoring of a lanthanide-binding peptide to a target protein enhances the paramagnetic anisotropic effect. Journal of biomolecular NMR 44, 157–166 (2009).

27. J. Jee, R. Ishima, A. M. Gronenborn, Characterization of specific protein association by 15N CPMG relaxation dispersion NMR: the GB1(A34F) monomer-dimer equilibrium. J Phys Chem B 112, 6008–6012 (2008).

28. J. Kuszewski, A. M. Gronenborn, G. M. Clore, Improving the packing and accuracy of NMR structures with a pseudopotential for the radius of gyration. Journal of the American Chemical Society 121, 2337–2338 (1999).

29. J. R. Lewandowski, M. E. Halse, M. Blackledge, L. Emsley, Protein dynamics. Direct observation of hierarchical protein dynamics. Science 348, 578–581 (2015).

30. J. A. Davey, R. A. Chica, Optimization of rotamers prior to template minimization improves stability predictions made by computational protein design. Protein Sci 24, 545–560 (2015).

31. J. K. Myers, C. N. Pace, J. M. Scholtz, Denaturant m values and heat capacity changes: relation to changes in accessible surface areas of protein unfolding. Protein Sci 4, 2138–2148 (1995).

32. I. R. Kleckner, M. P. Foster, An introduction to NMR-based approaches for measuring protein dynamics. Biochim Biophys Acta 1814, 942–968 (2011).

33. J. Jee, I. J. Byeon, J. M. Louis, A. M. Gronenborn, The point mutation A34F causes dimerization of GB1. Proteins 71, 1420–1431 (2008).

34. E. Campbell, M. Kaltenbach, G. J. Correy, P. D. Carr, B. T. Porebski, E. K. Livingstone, L. Afriat-Jurnou, A. M. Buckle, M. Weik, F. Hollfelder, N. Tokuriki, C. J. Jackson, The role of protein dynamics in the evolution of new enzyme function. Nature chemical biology 12, 944–950 (2016).

35. Y. Zhang, J. Skolnick, Scoring function for automated assessment of protein structure template quality. Proteins 57, 702–710 (2004).

36. J. Skolnick, D. Kihara, Y. Zhang, Development and large scale benchmark testing of the PROSPECTOR_3 threading algorithm. Proteins 56, 502–518 (2004).

37. T. Gallagher, P. Alexander, P. Bryan, G. L. Gilliland, 2 Crystal-Structures of the B1 Immunoglobulin-Binding Domain of Streptococcal Protein-G and Comparison with Nmr. Biochemistry. 33, 4721–4729 (1994).

38. Chemical Computing Group Inc. (Chemical Computing Group Inc., 1010 Sherbooke St. West, Suite #910, Montreal, QC, Canada, H3A 2R7, 2012).

39. P. Labute, Protonate3D: Assignment of ionization states and hydrogen coordinates to macromolecular structures. Proteins-Structure Function and Bioinformatics 75, 187–205 (2009).

40. I. W. Davis, W. B. Arendall 3rd, D. C. Richardson, J. S. Richardson, The backrub motion: how protein backbone shrugs when a sidechain dances. Structure 14, 265–274 (2006).

41. F. Lauck, C. A. Smith, G. F. Friedland, E. L. Humphris, T. Kortemme, RosettaBackrub-a web server for flexible backbone protein structure modeling and design. Nucleic Acids Research 38, W569–W575 (2010).

42. S. G. Nash, A survey of truncated-Newton methods. J. Comput. Appl. Math. 124, 45–59 (2000).

43. J. Wang, P. Cieplak, P. A. Kollman, How well does a restrained electrostatic potential (RESP) model perform in calculating conformational energies of organic and biological molecules? Journal of Computational Chemistry 21, 1049–1074 (2000).

44. R. A. Chica, M. M. Moore, B. D. Allen, S. L. Mayo, Generation of longer emission wavelength red fluorescent proteins using computationally designed libraries. Proc Natl Acad Sci USA 107, 20257–20262 (2010).

45. B. D. Allen, S. L. Mayo, Dramatic performance enhancements for the FASTER optimization algorithm. J Comput Chem 27, 1071–1075 (2006).

46. B. D. Allen, S. L. Mayo, An efficient algorithm for multistate protein design based on FASTER. J Comput Chem 31, 904–916 (2010).

47. R. L. Dunbrack, F. E. Cohen, Bayesian statistical analysis of protein side-chain rotamer preferences. Protein Science 6, 1661–1681 (1997).

48. S. L. Mayo, B. D. Olafson, W. A. Goddard, Dreiding - a Generic Force-Field for Molecular Simulations. J. Phys. Chem. 94, 8897–8909 (1990).

49. B. I. Dahiyat, S. L. Mayo, Probing the role of packing specificity in protein design. Proc Natl Acad Sci U S A 94, 10172–10177 (1997).

50. T. Lazaridis, M. Karplus, Discrimination of the native from misfolded protein models with an energy function including implicit solvation. Journal of Molecular Biology 288, 477–487 (1999).

51. E. K. Koepf, H. M. Petrassi, M. Sudol, J. W. Kelly, WW: An isolated three-stranded antiparallel beta-sheet domain that unfolds and refolds reversibly; evidence for a structured hydrophobic cluster in urea and GdnHCl and a disordered thermal unfolded state. Protein Science: A Publication of the Protein Society 8, 841–853 (1999).

52. F. Delaglio, S. Grzesiek, G. W. Vuister, G. Zhu, J. Pfeifer, A. Bax, NMRPipe: A multidimensional spectral processing system based on UNIX pipes. Journal of biomolecular NMR 6, 277–293 (1995).

53. B. A. Johnson, R. A. Blevins, NMR View: A computer program for the visualization and analysis of NMR data. Journal of biomolecular NMR 4, 603–614 (1994).

54. D. Wishart, B. Sykes, F. Richards, The chemical shift index: a fast and simple method for the assignment of protein secondary structure through NMR spectroscopy. Biochemistry 31, 1647–1651 (1992).

55. N. A. Farrow, O. Zhang, J. D. Forman-Kay, L. E. Kay, A heteronuclear correlation experiment for simultaneous determination of 15N longitudinal decay and chemical exchange rates of systems in slow equilibrium. Journal of biomolecular NMR 4, 727–734 (1994).

56. Y. Shen, F. Delaglio, G. Cornilescu, A. Bax, TALOS+: a hybrid method for predicting protein backbone torsion angles from NMR chemical shifts. Journal of biomolecular NMR 44, 213–223 (2009).

57. P. Güntert, in Protein NMR Techniques, A. K. Downing, Ed. (Humana Press, Totowa, NJ, 2004), pp. 353–378.

58. C. A. Smith, T. Kortemme, Predicting the tolerated sequences for proteins and protein interfaces using RosettaBackrub flexible backbone design. PloS one 6, e20451 (2011).

59. J. A. Davey, R. A. Chica, Multistate Computational Protein Design with Backbone Ensembles. Methods Mol Biol 1529, 161–179 (2017).

60. I. W. Davis, A. Leaver-Fay, V. B. Chen, J. N. Block, G. J. Kapral, X. Wang, L. W. Murray, W. B. Arendall 3rd, J. Snoeyink, J. S. Richardson, D. C. Richardson, MolProbity: all-atom contacts and structure validation for proteins and nucleic acids. Nucleic Acids Res 35, W375–383 (2007).

61. A. Bhattacharya, R. Tejero, G. T. Montelione, Evaluating protein structures determined by structural genomics consortia. Proteins: Structure, Function, and Bioinformatics 66, 778–795 (2007).

62. Y. J. Huang, R. Powers, G. T. Montelione, Protein NMR Recall, Precision, and F-measure Scores (RPF Scores): Structure Quality Assessment Measures Based on Information Retrieval Statistics. Journal of the American Chemical Society 127, 1665–1674 (2005).

